# Gene expression and *in situ* protein profiling of candidate SARS-CoV-2 receptors in human airway epithelial cells and lung tissue

**DOI:** 10.1101/2020.04.07.030742

**Authors:** Jennifer A. Aguiar, Benjamin J-M. Tremblay, Michael J. Mansfield, Owen Woody, Briallen Lobb, Arinjay Banerjee, Abiram Chandiramohan, Nicholas Tiessen, Anna Dvorkin-Gheva, Spencer Revill, Matthew S. Miller, Christopher Carlsten, Louise Organ, Chitra Joseph, Alison John, Paul Hanson, Bruce M. McManus, Gisli Jenkins, Karen Mossman, Kjetil Ask, Andrew C. Doxey, Jeremy A. Hirota

## Abstract

In December 2019, SARS-CoV-2 emerged causing the COVID-19 pandemic. SARS-CoV, the agent responsible for the 2003 SARS outbreak, utilizes ACE2 and TMPRSS2 host molecules for viral entry. ACE2 and TMPRSS2 have recently been implicated in SARS-CoV-2 viral infection. Additional host molecules including ADAM17, cathepsin L, CD147, and GRP78 may also function as receptors for SARS-CoV-2.

To determine the expression and *in situ* localization of candidate SARS-CoV-2 receptors in the respiratory mucosa, we analyzed gene expression datasets from airway epithelial cells of 515 healthy subjects, gene promoter activity analysis using the FANTOM5 dataset containing 120 distinct sample types, single cell RNA sequencing (scRNAseq) of 10 healthy subjects, immunoblots on multiple airway epithelial cell types, and immunohistochemistry on 98 human lung samples.

We demonstrate absent to low ACE2 promoter activity in a variety of lung epithelial cell samples and low *ACE2* gene expression in both microarray and scRNAseq datasets of epithelial cell populations. Consistent with gene expression, rare ACE2 protein expression was observed in the airway epithelium and alveoli of human lung. We present confirmatory evidence for the presence of TMPRSS2, CD147, and GRP78 protein *in vitro* in airway epithelial cells and confirm broad *in situ* protein expression of CD147 in the respiratory mucosa.

Collectively, our data suggest the presence of a mechanism dynamically regulating ACE2 expression in human lung, perhaps in periods of SARS-CoV-2 infection, and also suggest that alternate receptors for SARS-CoV-2 exist to facilitate initial host cell infection.

## INTRODUCTION

In 2003, the severe acute respiratory syndrome (SARS) outbreak caused by the SARS coronavirus (CoV) resulted in 8096 probable cases with 774 confirmed deaths[1, 2] In patients with SARS, deaths were attributed to acute respiratory distress associated with diffuse bilateral pneumonia and alveolar damage[3]. In December 2019, SARS-CoV-2 emerged causing the COVID-19 pandemic. SARS-CoV-2 is spreading at a much more rapid rate than SARS-CoV [4–6]. Similar clinical reports of diffuse bilateral pneumonia and alveolar damage have been reported[7–9]. Severe cases of SARS-CoV-2 have been associated with infections of the lower respiratory tract with detection of the virus throughout this tissue as well as the upper respiratory tract[7–9]. The biological mechanisms that may govern differences in the number of SARS and COVID-19 cases remain undefined. It is possible that SARS-CoV-2 possesses distinct molecular mechanisms that impact the virulence through viral proteins, greater susceptibility of host cells to infection, permissivity of host cells to virus replication, or some combination of these and other potentially unknown factors[10–13]. Understanding SARS and SARS-CoV-2 virus similarities and differences at the molecular level in the host may provide insights into transmission, pathogenesis, and interventions.

The seminal report identifying the receptor for SARS-CoV used a HEK293 cell over-expression system to identify angiotensin-converting enzyme 2 (ACE2) as a receptor by co-immunoprecipitation with SARS-CoV spike domain 1[14]. Subsequently, spike protein of SARS-CoV was identified as the viral interacting partner of ACE2. Host protease activity by TMPRSS2 facilitates ACE2 ectodomain cleavage and fusion of SARS-CoV membrane with host cell membrane[15–17]. ADAM17 has also been demonstrated to cleave ACE2 ectodomain, but this was not required for SARS CoV infection[18–20]. Mechanisms of SARS CoV entry distinct from ACE2 have also been reported and include activation by endosomal cathepsin L and cell surface expression of CD147 or GRP78[21–23] Each of these receptors were mechanistically interrogated and suggest that SARS CoV could initiate host cell entry and infection using multiple mechanisms. Recent *in vitro* reports have demonstrated that similar host proteins are involved in facilitating cell entry by SARS-CoV-2, such as ACE2 and TMPRSS2[5, 24] Biophysical and structural evidence strongly support an interaction of ACE2 with SARS-CoV-2 spike protein, similar to SARS-CoV spike protein[12, 13]. Molecular docking studies have also suggested that SARS-CoV-2 spike protein can interact with cell-surface GRP78[25]. Indirect evidence for a role of CD147 in SARS-CoV-2 binding has been demonstrated *in vitro* with the use of an anti-CD147 intervention that prevented virus replication[26]. Furthermore, a clinical study with an anti-CD147 intervention reduced symptoms and duration of hospital admission for COVID-19 patients[27]. In summary, although there is evidence that SARS-CoV-2 and SARS-CoV both utilize ACE2 as a receptor to facilitate virus entry, it is possible that differences in host-entry mechanisms play a role in the large epidemiological differences between the two viruses, which may include additional unidentified receptors.

ACE2 and TMPRSS2 were identified as cellular entry determinants for SARS-CoV using mechanistic studies. The original report of *in situ* human lung ACE2 expression described positive immunohistochemical staining for alveoli and airway epithelial cells, and immunocytochemical staining in A549 type II alveolar epithelial cells[28]. ACE2 protein expression is also present in the human lung adenocarcinoma cell line, Calu-3[29]. Similar to ACE2, the original report describing the expression of TMPRSS2 in human respiratory mucosa described expression in airway epithelium and type II alveolar epithelial cells[30]. The specificity of the ACE2 and TMPRSS2 antibodies used for analysis of expression patterns in human lung tissues remains to be addressed.

To address the uncertainties related to SARS-CoV-2 receptors in human lung, we performed gene expression and *in situ* protein profiling of candidate receptors in human airway epithelial cells and lung tissue. Our computational analysis used publicly available microarray gene expression datasets from airway epithelial cells of 515 unique subjects, single cell sequencing data from 10 subjects, and the FANTOM5 dataset for promoter activities of 74 lung-related cell and tissue types. For our *in situ* protein profiling, we performed immunohistochemical analysis of 98 human lung tissue samples. To determine antibody specificity, we performed immunoblots on protein isolated from Calu-3 cells, primary human airway epithelial cells, primary type II alveolar epithelial cells, the human bronchial epithelium cell (HBEC)-6KT cell line, the A549 type II alveolar epithelial cell line, and HEK cells. Collectively our data contrast previous reports, demonstrating rare ACE2 protein expression in the airway epithelium and alveoli of human lung. Our protein expression data are consistent with low ACE2 promoter activity in a variety of lung epithelial cell samples and low *ACE2* gene expression in both microarray and single cell RNA sequencing (scRNAseq) datasets. We present confirmatory evidence for the presence of TMPRSS2, CD147, and GRP78 protein *in vitro* in airway epithelial cells and confirm broad *in situ* protein expression of CD147 in the respiratory mucosa. Our data suggest that the presence of a mechanism dynamically regulating ACE2 expression in human lung, perhaps in periods of SARS-CoV-2 infection and/or that alternate receptors for SARS-CoV-2 exist to facilitate initial host cell infection in lung tissue.

## METHODS

### Human ethics

Procurement of primary human airway epithelial cells used for immunoblots and lung tissue for immunohistochemistry was approved by Hamilton integrated Research Ethics Board (HiREB 5099T, 5305T, 11-3559 and 13-523-C). UBC Research Ethics Office approved heart tissue archives and primary human airway epithelial cell collection.

### Upper and lower airway gene expression analysis

Public microarray experiments using Affymetrix chips (HuGene-1.0-st-v1 and HG-U133 Plus 2) on airway epithelial cell samples collected from nasal (GSE19190) or bronchial (GSE11906) brushings of healthy, non-smokers were obtained from the NCBI Gene Expression Omnibus (GEO) database [31, 32]. This resulted in a total of 80 individual samples from the two different experiments which includes 11 upper airway samples (Nasal: 11) and 69 lower airway samples (Trachea: 17, Large Airway: 17, Small Airway: 35). For all dataset samples, raw intensity values and annotation data were downloaded using the *GEOquery* R package (version 2.52.0)[33] from the Bioconductor project [34]. Probe definition files were downloaded from Bioconductor and probes were annotated using Bioconductor’s *annotate* package. All gene expression data were unified into a single dataset that was then RMA-normalized, and only genes present in both of the Affymetrix platforms (N = 16,013) were kept for subsequent analyses. Correction of experiment-specific batch effects was performed using the ComBat method[35] implemented using the *sva* R package (version 3.32.1)[36]. RMA-normalized expression levels for conventional (*ACE2, TMPRSS2, ADAM17*, and *CTSL*) and non-conventional (*CD147*, and *GRP78*) SARS-CoV-2 receptor genes were compared across the four defined airway levels, with *CDH1* expression level included as a positive control with known expression in lung tissue. Gene expression levels were tested for significant differences via pairwise Wilcoxon rank sum tests with Benjamini-Hochberg multiple testing correction using the *stats* R package (version 3.6.1). Gene expression box plots were generated with the *ggplot2* R package (version 3.2.1).

### Analysis of Curated Bronchial Epithelial Cell Brushing Dataset

A total of 1,859 public microarray experiments using Affymetrix chips (HG-U133 Plus 2 and HuGene-1.0-st-v1) on airway epithelial cell samples were selected from the NCBI GEO database. These samples were further filtered by removing individuals with asthma or COPD, resulting in a total of 504 individual healthy samples (GSE4302, 28 samples; GSE67472, 43 samples; GSE37147, 159 samples; GSE108134, 274 samples). Within this dataset, sex and age information was included for 310 samples with 86 females/106 males.

For all dataset samples, raw intensity values and annotation data were downloaded as described above. Probe definition files were retrieved as described above. All gene expression data were unified into a single dataset that was then RMA-normalized, and only genes present in both of the Affymetrix platforms (N = 16,105) were kept for subsequent analyses. Correction of experiment-specific batch effects was performed as described above.

### Analysis of promoter activity from the FANTOM5 dataset

The FANTOM5 promoterome dataset[37] for the hg38 assembly[38] was used to examine promoter activity of SARS-CoV-2-related human genes, namely *ACE2, TMPRSS2, ADAM17, CTSL* (cathepsin L1), CD147 and GRP78. Using the ZENBU genome browser[39], the nearest cap analysis of gene expression (CAGE) peak upstream and on the same strand as each of the aforementioned genes was extracted and analyzed. The dataset consists of CAGE promoter activity data for 1,886 primary cells, cell lines, and tissues from humans, and quantified as normalized transcripts per million (TPM). A subset of FANTOM5 CAGE data (120 samples) is presented considering only samples related to lung, gut, heart, and prostate tissues (consisting of 74, 19, 15, and 12 samples, respectively). Normalized TPM values for each CAGE peak, an approximation for promoter activity, were log_10_ transformed and separated according to tissue and cell type, and the radius of each point is proportional to these transformed normalized TPM values.

### Analysis of single cell RNA sequencing (scRNAseq) data

Data preprocessed using the Cell Ranger pipeline (10x Genomics) were obtained from GSE135893. Samples from 10 control subjects and 12 IPF patients were downloaded and post-processed with *Seurat* package in R[40]. Cell populations were defined using the markers provided in the source paper[41]. Cells belonging to the 10 control subjects were used for further analysis. Visualizations of violin plots were created using *Seurat*.

### Primary Human Airway Epithelial Cells

The human lung adenocarcinoma cell line, Calu-3, was grown under culture conditions defined by the supplier (ATCC-HTB-55). Primary human airway epithelial cells isolated via bronchial brushings from consented healthy individuals were grown in PneumaCult ExPlus (Stemcell Technologies, Vancouver Canada) under submerged monolayer culture conditions and used in between passage 1 and 4. The human bronchial epithelial cell line, HBEC-6KT, was grown under submerged monolayer culture conditions in keratinocyte serum free media supplemented with epidermal growth factor (0.4ng/ml) and bovine pituitary extract (50μg/ml)[42–45].

### Immunoblots

Cell protein was isolated using RIPA lysis buffer (VWR, Ontario, Canada) supplemented with protease inhibitor cocktail (Sigma, Ontario, Canada) with quantification performed using Bradford assay reagents (Bio Rad, Ontario, Canada). Immunoblots were performed using stain free 4-20% pre-cast gradient gels and imaged on a ChemiDoc XRS+ Imaging system (Bio Rad, Ontario, Canada). For each immunoblot, 20μg of protein was added per lane. ACE2 (R&D Systems - MAB933 – Monoclonal - Clone 171606 – 2μg/ml), TMPRSS2 (Atlas Antibodies - HPA035787 – Polyclonal – 0.4μg/ml), CD147 (Abcam – ab666 – Monoclonal – Clone MEM-M6/1 – 1μg/ml), and GRP78 (BD – 610979 – Monoclonal – Clone – 40/BiP – 0.25μg/ml) primary antibodies were diluted in 5% skim milk/Tris buffered saline with 0.1% Tween-20 and incubated overnight on a rocker at 4°C with detection performed the following day using an anti-mouse-HRP (ACE2, CD147, GRP78) or anti-rabbit-HRP (TMPRSS2) conjugated secondary antibodies at 1:3000 for 2hrs at room temperature (Cell Signaling, Danvers, MA, USA). Visualization of TMPRSS2, CD147, and GRP78 was performed using was performed using Clarity™ Western enhanced chemiluminescence (ECL) Substrate, while ACE2 was visualized with Clarity Max™ ECL Substrate (Bio Rad, Ontario Canada). Total protein loading images were collected as a confirmation of equal protein loading between sample types[46]. The immunogen for ACE2 primary antibody is mouse myeloma cell line NS0-derived recombinant human ACE2 Gln18-Ser740 (predicted). The immunogen for TMPRSS2 primary antibody is the recombinant protein epitope signature tag antigen sequence, GSPPAIGPYYENHGYQPENPYPAQPTVVPTVYEVHPAQYYPSPVPQYAPRVLTQASNPVVCTQPKSPSGTVCTSKT. The immunogen for the CD147 primary antibody is recombinant full-length protein corresponding to human CD147. The immunogen for the GRP78 primary antibody is human BiP/GRP78 amino acids 525-628.

Independent immunoblot analysis (L.O, C.J. A.J. and G.J) were performed on A549, HEK, and immortalized human bronchial epithelial cells. Equal amounts of protein (20µg) were loaded on to 4-12%, Bis-Tris gradient gels (ThermoFisher, NP0326BOX) with anti-ACE2 (Abcam – ab108252 – Rabbit monoclonal – Clone EPR4435(2) – 1/500 dilution of stock antibody). A loading control of GAPDH was used to demonstrate protein loading (Abcam – ab181603 – Rabbit monoclonal – EPR16884 – 1/10000 dilution of stock antibody). Visualization was performed with ECL Clarity (Biorad, UK) on a Licor C-DiGit.

### Immunohistochemistry

Formalin-fixed paraffin-embedded human lung tissue from non-diseased regions were obtained from archived tissue blocks from patients who had undergone lung resection for clinical care. Human heart tissue was from the UBC Cardiovascular Tissue Registry. Four-micron thick sections were cut and stained for ACE2 (15μg/ml), TMPRSS2 (10μg/ml), and CD147 (5μg/ml) using the same antibodies used for immunoblot analysis. All staining was performed on a Leica Bond RX system with Leica Bond reagents, heat-induced antigen retrieval in at pH 6 (20 minutes) with primary antibody incubation for 20 minutes. Digital slide scanning was performed using an Olympus VS120-L100 Virtual Slide System at 40X magnification with VS-ASW-L100 V2.9 software and a VC50 colour camera followed by image visualization with HALO image analysis software.

## RESULTS

### Candidate genes important in SARS-CoV-2 infection are detectable at varying levels in human airway epithelial cells and lung tissue

We performed a targeted analysis of *ACE2, TMPRSS2, ADAM17, CTSL, CD147*, and *GRP78* gene expression as candidates important for SARS-CoV-2 infection in human airway epithelial cells. Here and throughout gene expression analyses, *CDH1* (E-cadherin) was used as a control for lung epithelial cell phenotype. We first examined these genes in a curated dataset of upper and lower airway epithelial cell gene expression from the nasal sinus to the 12^th^ generation of airway in the lung (**Figure 1A**).

**Figure 1.**
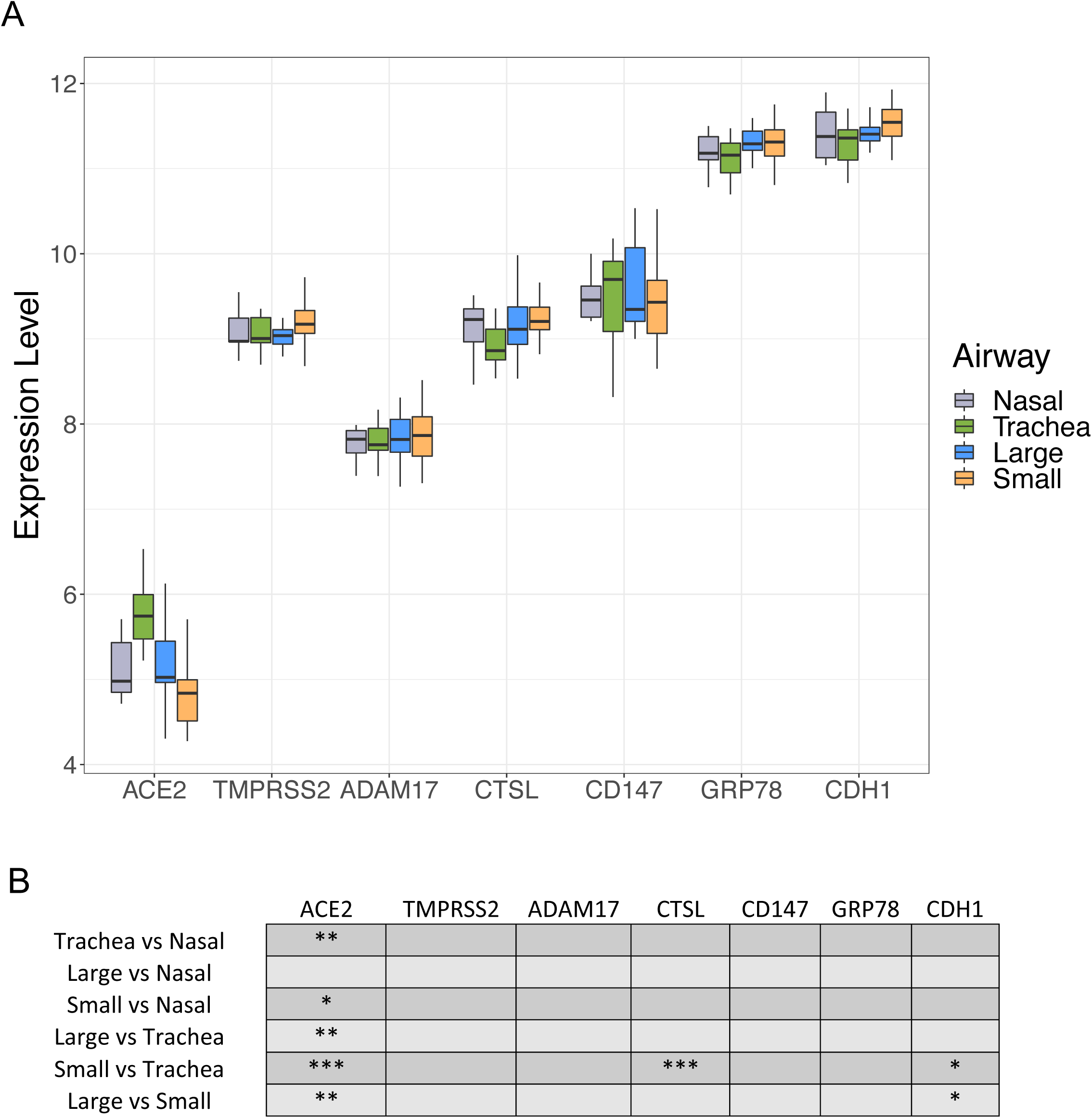
Microarray expression profiles of candidate SARS-CoV-2 receptor genes in upper and lower airways. **A:** Normalized log_2_ expression levels for *ACE2, TMPRSS2, ADAM17, CTSL, CD147*, and *GRP78* genes compared across the upper airway (nasal, grey) and lower airways (trachea, green; large airway, blue; small airway, orange). *CDH1* gene expression level is included as a positive control. **B:** Statistical values for comparisons for each gene at each airway generation. *=*p*<0.05; **=*p*<0.01; ***=*p*<0.001.

In the upper airways, all candidates were expressed with the highest levels observed for *GRP78* and the lowest level observed for *ACE2*. Analysis along multiple generations of the lower airways (trachea, large (4-6^th^ generation), and small airways (10-12^th^ generation) revealed identical relative expression patterns with *ACE2* being the least expressed and *GRP78* being the highest expressed. *ACE2* gene expression showed the greatest variability along the upper and lower airways, with greatest expression observed in the trachea samples and the lowest expression in the small airway (**Figure 1B**).

Following our observation of consistent expression along the upper and lower airways of candidate genes important in SARS-CoV, we determined if sex or age impacted gene expression levels in healthy individuals using a curated dataset of bronchial brushings from 504 healthy subjects (**Table 1**). The expression levels for the candidate genes in healthy subjects paralleled the patterns observed in the smaller survey of upper airways, trachea, large, and small airways (**Figure 2**). Median *ACE2* gene expression was the lowest, while *GRP78* gene expression was the highest (**Figure 2A**). No gene candidate demonstrated sex dependence for expression levels (**Figure 2B**). No microarray chip dependent effects were observed for relationships between sex and gene expression. For quantitative analyses related to age and gene expression our curated database was divided into datasets that used either the HG-U133 Plus 2 or HuGene-1.0-st-v1 microarray due to differences in age distributions. In the HuGene-1.0-st-v1dataset (n=181) which included a greater proportion of older (>50) individuals, we observed reduced *ACE2* gene expression with age (**Figure 2C**, *p* < 0.05).

**Figure 2:**
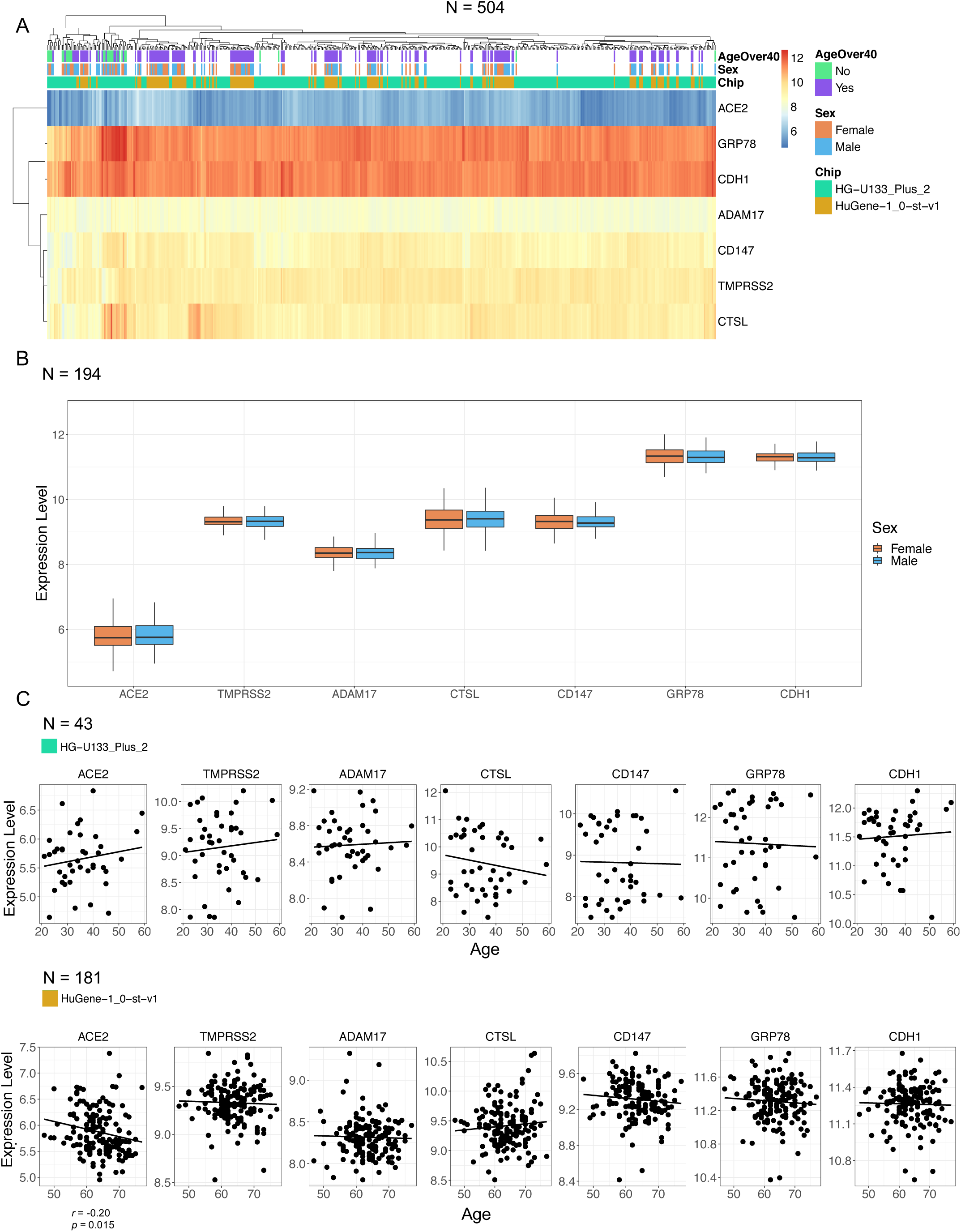
Microarray expression profiles of candidate SARS-CoV-2 receptor genes in lower airway epithelial cells analyzed by age and sex. **A:** Clustered heatmap of log_2_ expression levels from 504 GEO samples, annotated by age, sex, and microarray chip platform. Expression values reflect signal intensities, indicating lowest detected expression of *ACE2* and highest expression of *GRP78* and *CDH1*. **B:** Per-gene boxplots of expression levels separated by sex. **C:** Plots of gene expression levels versus age, with linear regression lines of best fit. A weak negative correlation (*r* = −0.20, *p* = 0.015) was detected for *ACE2* in the second dataset. Correlations were performed separately between platforms because of differences in their age distributions.

Promoter activity data of each of the candidate genes important in SARS-CoV-2 binding and infection were extracted and analyzed from the FANTOM5 dataset, which includes 1,886 primary cells, cell lines, and tissue sample types (**Figure 3**). We selected all samples formats that included “lung”, “nasal”, “airway”, “olfactory” to identify lung-specific sample types. Gut, heart, and prostate tissue samples were analyzed as controls. Consistent with our observed gene expression analysis along the upper and lower airways, normalized TPM values for each CAGE peak demonstrate that *CD147* promoter activity was elevated relative to *ACE2* promoter activity across airway epithelial cells and lung tissue samples. *Cathepsin L* promoter activity was the lowest of all candidate genes, which contrasted the modest expression observed at the gene level (**Figure 2A**). Both microarray gene expression analysis and promoter activity were consistent with results of candidate gene expression in a scRNAseq dataset of 10 healthy subjects (**Supplementary Figure 1**).

**Figure 3:**
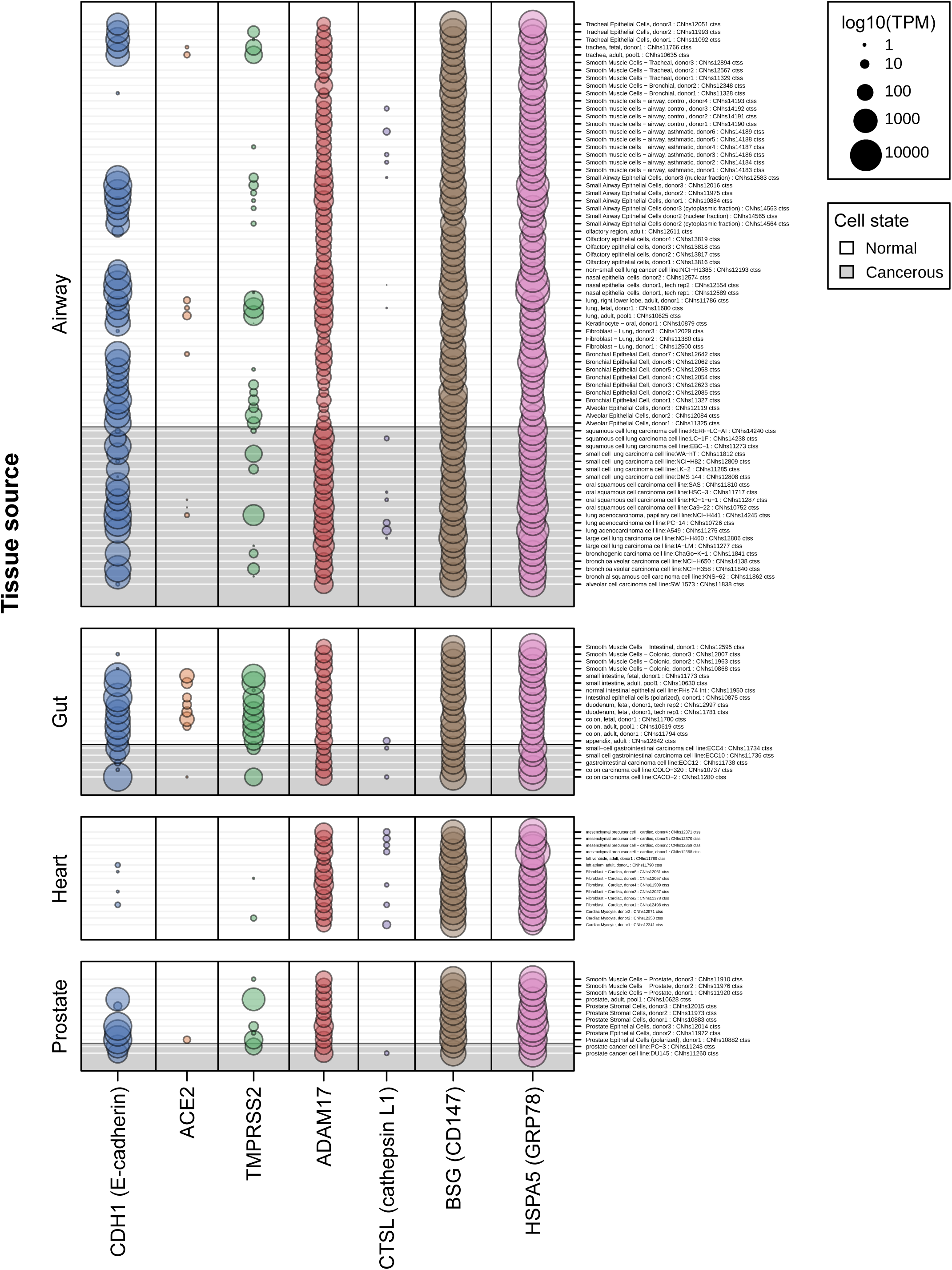
CAGE promoter activity for SARS-CoV-2-related genes from the FANTOM5 CAGE dataset. The FANTOM5 CAGE data consists of quantified promoter expression levels across the human genome for 1,866 samples from primary cells, cell lines, and tissue samples. In this figure, the FANTOM5 CAGE promoter activity data for several SARS-CoV-2-related genes are shown for samples related to lung, gut, heart, and prostate tissues (n=120). Dot sizes are proportional to promoter activity, depicted as log_10_-transformed normalized transcripts per million (TPM). Notably, ACE2 is either not expressed or expressed at low levels (less than 1 TPM in all but one sample) in the airway, including measurements from healthy (white rows) and cancerous cells (grey rows).

Collectively, our gene expression analysis of the upper and lower airways of healthy males and females of diverse ages suggests that *ACE2* gene expression is low relative to all other candidate SARS-CoV-2 receptor genes analyzed in human airway epithelial cells. Furthermore, we observe no sex-dependent or age-dependent expression patterns of any candidates at the gene level.

### *In vitro* and *in situ* protein profiling reveals distinct expression patterns for candidates important in SARS-CoV-2 infection

Analysis of transcriptional data may not be indicative of *in situ* protein expression levels[47]. To extend our gene expression observations, we performed *in vitro* immunoblots on human airway epithelial cell lysates and *in situ* protein immunohistochemistry on human lung tissue using the same antibodies for each method.

An anti-ACE2 antibody detected only a single band in Calu-3 cells at the predicted molecular weight (∼110kDa) of ACE2 protein. The anti-ACE2 antibody required the use of a super-sensitive ECL solution (**Figure 4A** – Top panel - Lanes 1-3). No ACE2 protein was detected in primary airway epithelial cells or the HBEC-6KT cell line despite confirmation of protein loading (**Figure 4A** – Lanes 4-9, bottom panel confirms protein loading). Independent immunoblotting with a distinct anti-ACE2 primary antibody was performed, with a single band observed in HEK cells, but not in immortalized human bronchial epithelial cells or A549 cells (**Supplement Figure 2**).

**Figure 4:**
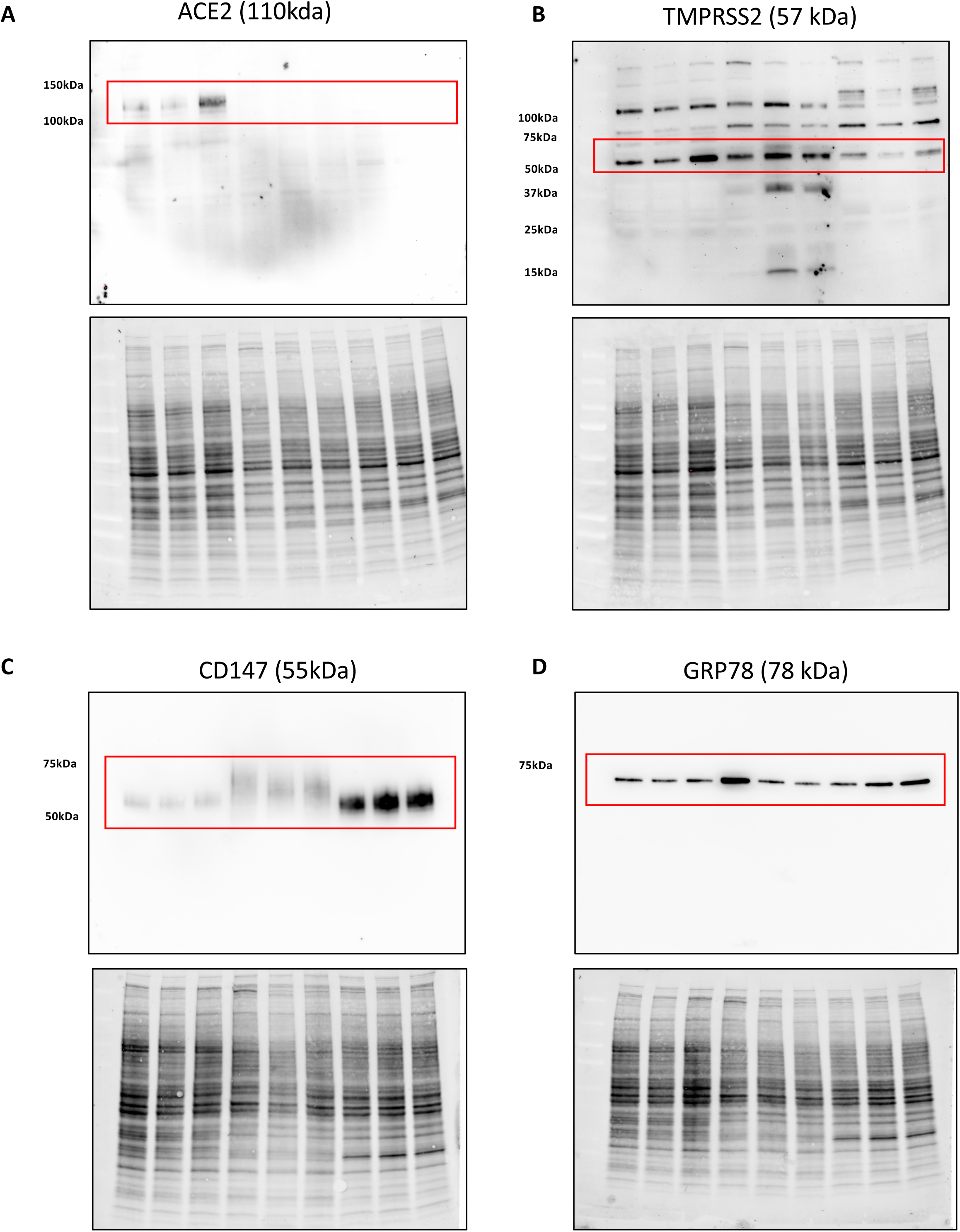
Immunoblot analysis of ACE2, TMPRSS2, CD147, and GRP78 protein expression in human airway epithelial cell protein lysates. **A:** ACE2 with single band for predicted molecular weight of 110kDa. **B:** TMPRSS2 with multiple bands including a dominant band at predicted molecular weight of 57 kDa. **C:** CD147 with a single broad band around predicted molecular weight of 55kDa. **D:** GRP78 with a single band at predicted molecular weight of 78 kDa. Lanes 1-3: Calu-3 cells. Lanes 4-6: Primary human airway epithelial cells. Lanes 7-9: HBEC-6KT. All cells grown under submerged monolayer conditions, with n=3 independent passages (Calu-3 or HBEC-6KT) or donor samples (Primary human airway epithelial cells – non-smoker, healthy subjects). For each independent blot of each protein, all of the same samples were run. Total protein loading control provided to demonstrate equal protein loaded for each sample.

An anti-TMPRSS2 antibody detected multiple bands in all airway epithelial cell samples with a dominant band at the predicted molecular weight of ∼55kDa (**Figure 4B** – Top panel). These patterns were conserved across all cell types that were analyzed.

An anti-CD147 antibody detected a single band in all airway epithelial cell samples with a dominant band at the predicted molecular weight of ∼55kDa (**Figure 4C –** Top panel). The immunoblot bands were consistent with the heavy glycosylation of CD147[48].

An anti-GRP78 antibody detected a single band in all airway epithelial cell samples with a dominant band at the predicted molecular weight of ∼78kDa (**Figure 4D –** Top panel).

The immunoblots using anti-ACE2, CD147, and GRP78 demonstrated a single band of predicted molecular weight, suggesting that observed immunohistochemical staining should be specific to the protein of interest based on the target epitope, as both methods detect denatured proteins[49]. The same anti-ACE2 and CD147 antibodies were validated for immunohistochemistry. The anti-TMPRSS2 was used for immunohistochemistry, although the multiple bands observed by immunoblot caution the specificity of any observed *in situ* staining. Attempts to optimize anti-GRP78 antibody application for immunohistochemistry were unsuccessful.

ACE2 immunohistochemistry revealed only select staining in rare cells in the airways and the alveoli of all 98 human lung samples analyzed that included healthy subjects and those with chronic lung diseases (**Figure 5**). A single healthy human sample contained one positive airway epithelial cell with additional positive staining in the peripheral lung in cells with type II alveolar epithelial cell morphology (**Figure 5A – second row**). A representative image of a sample from a smoker with chronic obstructive pulmonary disease (**Figure 5B – second row**) shows no ACE2 protein staining in the airway epithelium and a rare positive cell in sub-basement membrane tissue. Lung microvasculature and human heart tissue stained had positive staining (**Supplementary Figure 3-4**) consistent with previously described reports for ACE2 protein staining patterns [50, 51].

**Figure 5:**
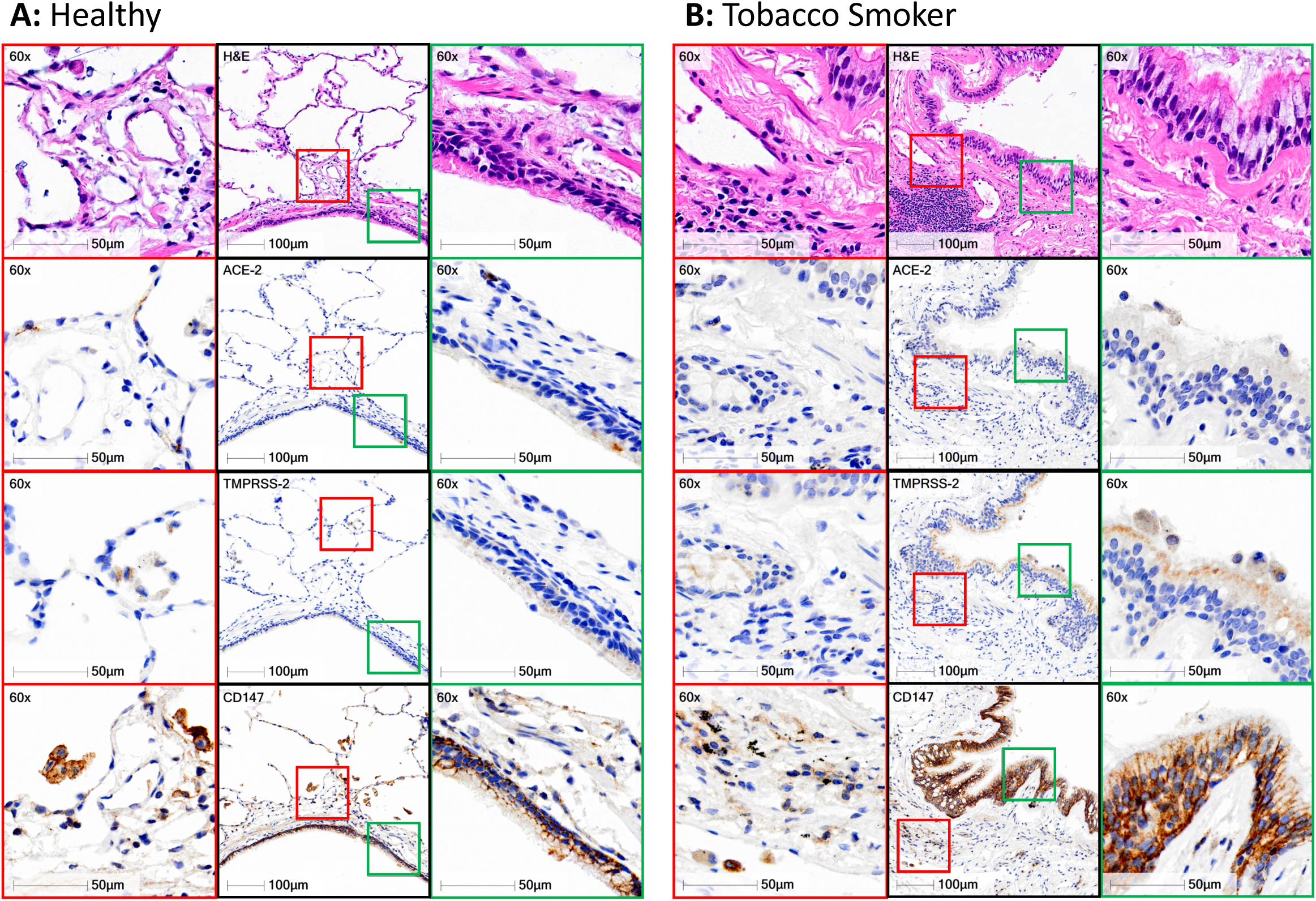
Immunohistochemical localization of ACE2, TMPRSS2, and CD147 protein in human lung tissue. **A:** Representative panel of a healthy non-smoker with no underlying chronic airway disease. **B:** Representative panel of a smoker with chronic obstructive pulmonary disease. For both panels, black squares represent low magnification (15X) of a conducting airway with airway epithelium. Green squares correspond to high magnification regions (60X) of conducting airway epithelium that are defined in the low magnification image. Red squares correspond to high magnification regions (60X) of lung tissue away from airway lumen that are defined in the low magnification image. Row 1 - hematoxylin & eosin, Row 2 – ACE2, Row 3 – TMPRSS2, Row 4 – CD147. Positive immunohistochemical staining is rust/brown. Total number of independent samples analyzes was 98.

TMPRSS2 immunohistochemistry revealed diffuse staining in the airway epithelium and in immune cells in the lung periphery, with greater staining in smokers with chronic obstructive pulmonary disease (COPD) (**Figure 5A-B – third row**). These observations were consistent in the majority of the 98 human samples examined.

CD147 immunohistochemistry revealed strong membrane restricted staining in the airway epithelium and diffuse staining in immune cells in the lung periphery, with greater staining in smokers with COPD (**Figure 5A-B – fourth row**). These observations were consistent in the majority of the 98 human samples examined.

Collectively, our *in vitro* and *in situ* protein profiling is consistent with our gene expression analysis with CD147 protein expression dominant over TMPRSS2 and ACE2. ACE2 protein expression is rare in human lung tissue and found in select cells in both healthy individuals and those with chronic lung diseases. TMPRSS2 and CD147 protein expression are potentiated in individuals with a history of tobacco smoking and a diagnosis of COPD.

## DISCUSSION

The global COVID-19 pandemic that emerged in late 2019 is caused by SARS-CoV-2. The possible host receptor(s) for SARS-CoV-2 have not been exhaustively surveyed in human lung tissue at the gene and protein level. Understanding the expression levels and localization of candidate SARS-CoV-2 receptors in host lung tissue may provide insights into therapeutic interventions that might reduce disease spread, viral replication, or disease pathology. To address this knowledge gap, we performed gene expression and *in situ* protein profiling of candidate receptors in human airway epithelial cells and lung tissue (**summarized in Figure 6**). Collectively our data demonstrate rare ACE2 protein expression in human airway epithelial cells *in vitro* and *in situ*. Our protein expression data are consistent with low ACE2 promoter activity in a panel of lung epithelial cell samples and low *ACE2* gene expression in bronchial epithelial cells (microarray) and lung cells (scRNAseq). We present confirmatory evidence for the presence of TMPRSS2, CD147, and GRP78 protein *in vitro* in airway epithelial cells and confirm broad *in situ* protein expression of CD147 in the respiratory mucosa. Our data suggest that for ACE2 to be an integral receptor for SARS-CoV-2, mechanisms are likely to exist that dynamically regulate expression in human lung, perhaps in periods of SARS-CoV-2 infection[52]. It is also possible that alternate receptors for SARS-CoV-2 are important in initial host cell infection.

**Figure 6:**
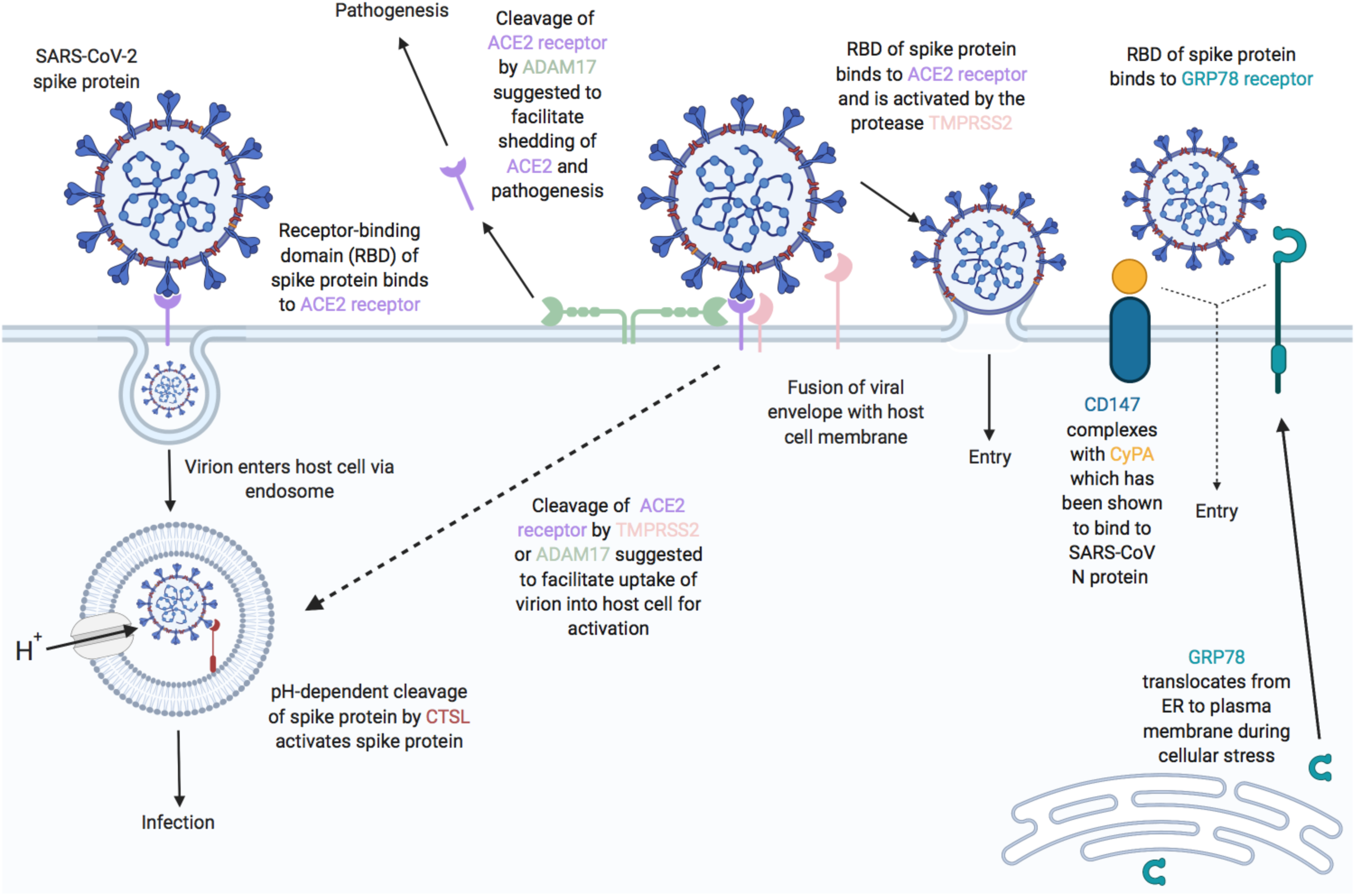
Proposed functions of host airway epithelial cell molecules for interaction with SARS-CoV-2. Proteins associated (or suggested to be associated) with host cell entry of SARS-CoV-2 and the activation of the SARS-CoV-2 spike protein (SARS-S) are displayed. ACE2 is suggested as the primary SARS-S receptor for viral entry (interaction of ACE2 receptor-binding domain (RBD) with SARS-S leading to endosomal viral uptake) followed by activation of SARS-S via pH-dependent CTSL-mediated cleavage. Secondary methods of viral entry and SARS-S activation are likely to involve proteases (e.g. TMPRSS2 and ADAM17) and/or secondary receptors (CD147 and GRP78). Dashed lines indicate mechanisms that have not been fully validated. Figure adapted from [20] with updates and additional information on candidate host molecules. Created with BioRender.com.

Using a curated microarray gene expression dataset generated from bronchial brushings of 504 healthy subjects that considers the limitations of merging multiple datasets from distinct sources, we observed that sex did not correlate with gene expression of any candidate host molecule involved in SARS-CoV-2 infection and that *ACE2* and *TMPRSS2* were the lowest expressed genes of interest examined. In one dataset, *ACE2* gene expression modestly decreased with age, although protein level confirmation was not possible. The low level of *ACE2* and *TMPRSS2* gene expression in bulk bronchial epithelial cell gene expression samples suggests low levels of cells expressing both of these genes within this lung tissue.

Advances in transcriptomics have enabled scRNAseq that has identified unique and rare cell types in human lung that may have importance in health and disease[53, 54]. scRNAseq provides an opportunity to look at transcriptional profiles in subsets of cell populations, which may isolate a cell signal from a bulk sample. We therefore utilized scRNAseq data from healthy human lung samples as a parallel approach. The resolution of scRNAseq for subpopulations of epithelial cells revealed low or absent expression of *ACE2* gene in all populations examined, whereas *CD147* and *GRP78* were present in all populations. Our results are consistent with current publicly available data that discuss the presence of a rare ACE2/TMPRSS2 positive cells[52]. Using lung samples from eight individuals (four HIV and active tuberculosis double +ve, two HIV +ve and tuberculosis -ve, and two double -ve controls), Ziegler et al. [52] have reported in humans that only 0.8% of type II alveolar epithelial cells expressed both *ACE2* and *TMPRSS2* genes. Further analysis of ciliated cells found that 5.3% of these cells expressed both *ACE2* and *TMPRSS2* genes. *In vitro* models with SARS-CoV, are consistent with this finding as ciliated cells are preferentially targeted by this coronavirus[55]. Most intriguing is that *ACE2* and *TMPRSS2* gene expressing cells were only identified in the HIV and tuberculosis double +ve samples. These observations were replicated in the upper airways, with only a rare population of secretory epithelial cells (0.3% of this population) co-expressing *ACE2* and *TMPRSS2*. The reported scRNAseq results are consistent with a focused analysis looking at only *ACE2* gene expression in a variety of lung cell types[56]. Importantly, these elegant transcriptomic analyses confirm our observations in bulk tissue microarray datasets.

Consortium based publicly available datasets represent another parallel approach to confirm our data. We have used the FANTOM5 dataset containing CAGE promoter activation data for 1,866 primary cells, cell lines, and tissue samples from humans[37] to examine the level of promoter activity for each candidate SARS-CoV-2-receptor genes. The FANTOM5 CAGE data provides an additional and complementary approach to quantifying gene expression since a given gene’s shared promoter can yield multiple transcripts at different expression levels, as well as being partially independent of any given transcript’s half-life in the cell. In general, the promoter activity of ACE2 in airway-related tissues is low or absent; only a single sample originating from an adult lung yields a normalized CAGE promoter expression level above one transcript per million, while expression was observed in gut cells, consistent with known patterns of ACE2 expression[57]. Consistent with the microarray data, CD147 promoter activity is elevated relative to ACE2 across airway-related cells and tissues, although the relatively low CTSL (cathepsin L1) promoter activity is incongruent with modest levels of gene expression.

The expression of genes does not always correlate with protein expression[47]. With this in mind we performed immunoblot analyses on three distinct airway epithelial cell sample types. We used the human Calu-3 adenocarcinoma cell line as this cell is susceptible and permissive to SARS-CoV-2 infection and expresses ACE2, an observation we confirm[24, 29]. We also used primary human airway epithelial cells and the bronchial epithelial cell line (HBEC-6KT). We performed immunoblots for ACE2 and TMPRSS2 as these have been highlighted as interacting with SARS-CoV-2, while we probed CD147 as recent pre-clinical and clinical studies have provided proof of concept as a candidate SARS-CoV-2 receptor[26, 27]. Lastly, GRP78 was dominantly expressed throughout transcriptomic studies and was selected as a positive control as previous expression has been confirmed in human airway epithelial cells[58]. Cathepsin L was excluded from the present analysis due to low promoter activity (**Figure 3**), while ADAM17 was excluded as the proposed function in coronavirus infections is via ACE2[5, 24], which was included in analysis. Immunoblot analysis with all antibodies revealing dominant bands of predicted molecular weight, with the anti-TMPRSS2 polyclonal antibody revealing additional minor bands in all cell samples examined. The identity of these other bands remains unclear and suggest downstream immunohistochemical analysis may be confounded by the specificity of this antibody. In contrast, antibodies for ACE2, CD147, and GRP78 were specific and could be used for immunohistochemistry without concerns of specificity. Interestingly, ACE2 protein could only be detected with a supersensitive ECL solution and only in Calu-3 cells, suggesting absent protein in primary human airway epithelial cell and the HBEC-6KT cell lines. Our data are consistent with previous immunoblots of primary human airway epithelial cells grown under submerged monolayer conditions using the same primary antibody, where ACE2 protein was absent, and only expressed under air-liquid interface culture conditions[59]. The observation that CD147 and GRP78 are also expressed in Calu-3 cells encourages further interrogation into these host proteins, as they may contribute to function of ACE2 and TMPRSS2 in SARS-CoV-2 binding and fusion in this cell type. Collectively, the profiling of antibodies by immunoblot of airway epithelial cells revealed distinct band patterns demonstrative of antibody specificity for ACE2, CD147, and GRP78, and to a lesser extent for TMPRSS2.

Immunohistochemical analysis has been performed for localization of ACE2 and TMPRSS2 in human lung[28, 30]. The observation of positive staining in human lung tissue for these proteins was not accompanied by companion immunoblot or complementary approaches to define the specificity of the antibody used[49]. In the absence of determination of antibody specificity, the historical data presented should be interpreted with caution. To address the issue of antibody specificity for immunohistochemical staining, we used the same antibodies we validated by immunoblot. We again focused on ACE2 and TMPRSS2 as these are candidate proteins important for SARS-CoV-2 infection of host cells. We performed parallel analysis of CD147, but not GRP78, as the latter antibody was not validated for immunohistochemistry at varying experimental conditions. Our immunohistochemical staining patterns of ACE2 were consistent with transcriptional profiling and immunoblots with only 1 of 98 human samples demonstrating rare staining in the airway and alveolar epithelium. Positive ACE2 staining in heart tissues and areas of lung microvasculature suggest our staining protocol was successful. These results directly contrast those reported with antibodies that lacked validation for specificity[28, 30]. TMPRSS2 was expressed more frequently across all samples examined with variability in the airway epithelium, associated with history of smoking and/or chronic obstructive pulmonary disease status. In contrast, CD147 expression was observed in airway epithelium of all samples. Similar to TMPRSS2, elevated CD147 expression was associated with history of smoking and/or chronic obstructive pulmonary disease status, consistent with previous reports[48].

Our study has several limitations that have not already been addressed above. Our observation of differences in gene expression between upper and lower airways and along the airway tree were not corroborated at the protein level. It remains possible that entirely different protein expression profiles for the candidate molecules examined exist in the upper airway, presenting a different environment for SARS-CoV-2 interaction with the respiratory mucosa. Nasal pharyngeal swabs are capable of detecting SARS-CoV-2 virus[60] and this anatomical region likely is important for subsequent infection in the lower airways[8]. Related to this potential temporality of effect, it is possible that SARS-CoV-2 induces the expression of receptors on host cells following infection[52]. Our study is also limited by examining candidate molecules important in SARS-CoV-2 infection under basal conditions, in the absence of viral or environmental stimuli which may regulate gene transcription and protein translation.

SARS-CoV-2 infection and transmission has caused the global COVID-19 pandemic. An understanding of the receptors used by SARS-CoV-2 for host cell infection and the parallel characterization in human samples is required to inform development of intervention strategies aimed at mitigating COVID-19. Our data demonstrate rare ACE2 protein expression in human airway epithelial cells *in vitro* and *in situ*, consistent with low ACE2 promoter activity and *ACE2* gene expression in bronchial epithelial cells. We present confirmatory evidence for the presence of TMPRSS2, CD147, and GRP78 protein *in vitro* in airway epithelial cells and confirm broad *in situ* protein expression of CD147 in the respiratory mucosa. Due to the overwhelming evidence that SARS virus interacts with ACE2, there are likely to be alternate mechanisms regulating ACE2 in the respiratory mucosa in the context of SARS-CoV-2 infection, and/or, perhaps other co-receptors, beyond what is expressed under basal conditions at the protein level.

## ACKNOWLEDGEMENTS

We would like to acknowledge Mary Jo Smith from the McMaster Immunology Research Centre Core Histology facility for her timely and professional expertise with antibody staining for immunohistochemistry. We would like to thank Drs. Sam Wadsworth and John McDonough for their intellectual discussions around CD147 and airway epithelial cell biology. We would like to thank Dr. Charles Plessy for his suggestion to analyze the FANTOM5 data. We would like to thank all of the personal and professional support from the author’s respective families and research institutes, and most importantly, the frontline healthcare workers during the COVID-19 pandemic.

## FIGURE LEGENDS

**Supplement Figure 1:**
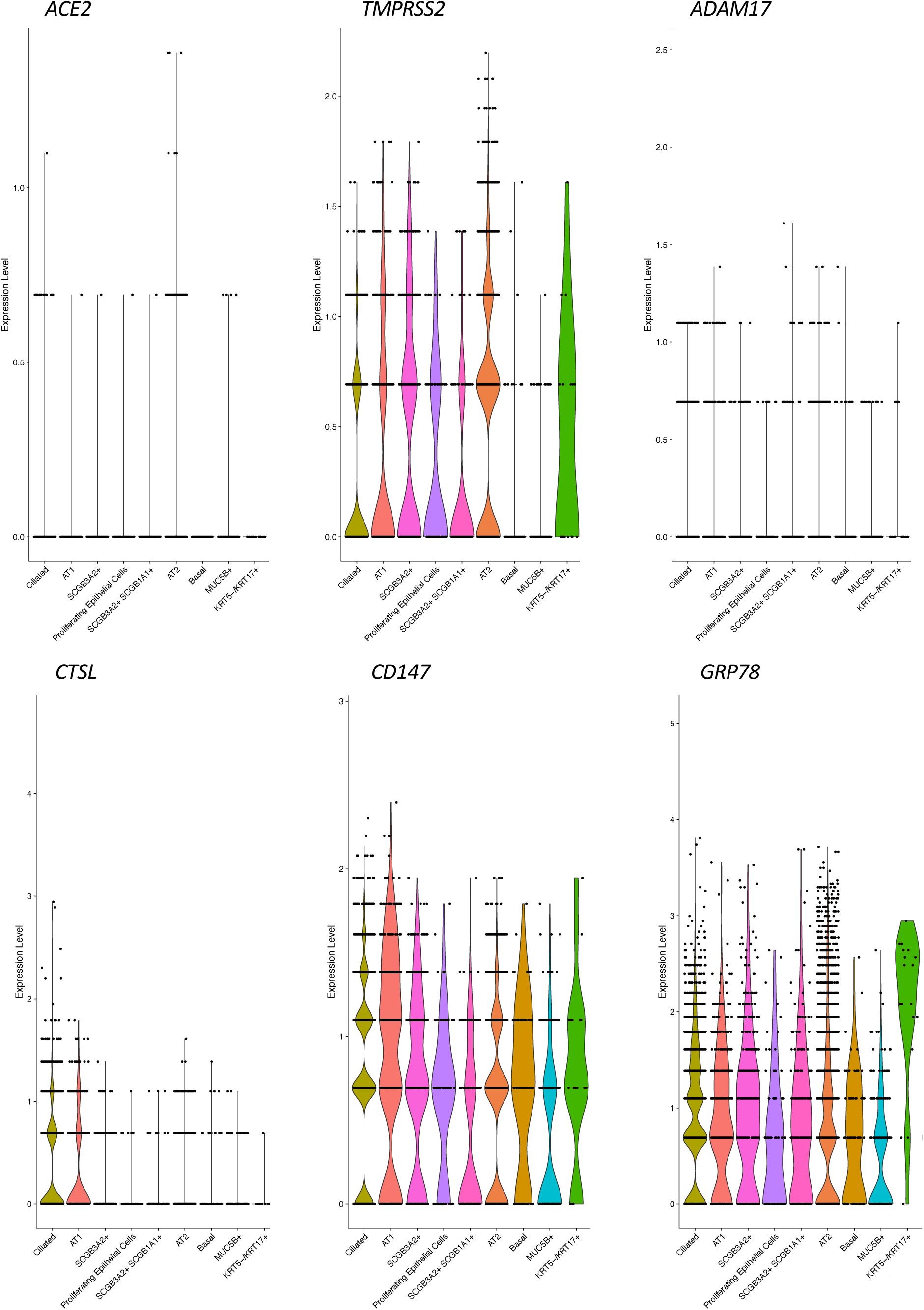
scRNAseq of peripheral lung tissue in nonfibrotic individuals. Violin plots for expression levels of **A:** *ACE2*, **B:** *TMPRSS2*, **C:** *ADAM17*, **D:** *CTSL (Cathepsin L)*, **E:** *CD147*, and **F:** *GRP78* across lung epithelial cell populations in healthy subjects (see methods for dataset reference for cell population markers).

**Supplement Figure 2:**
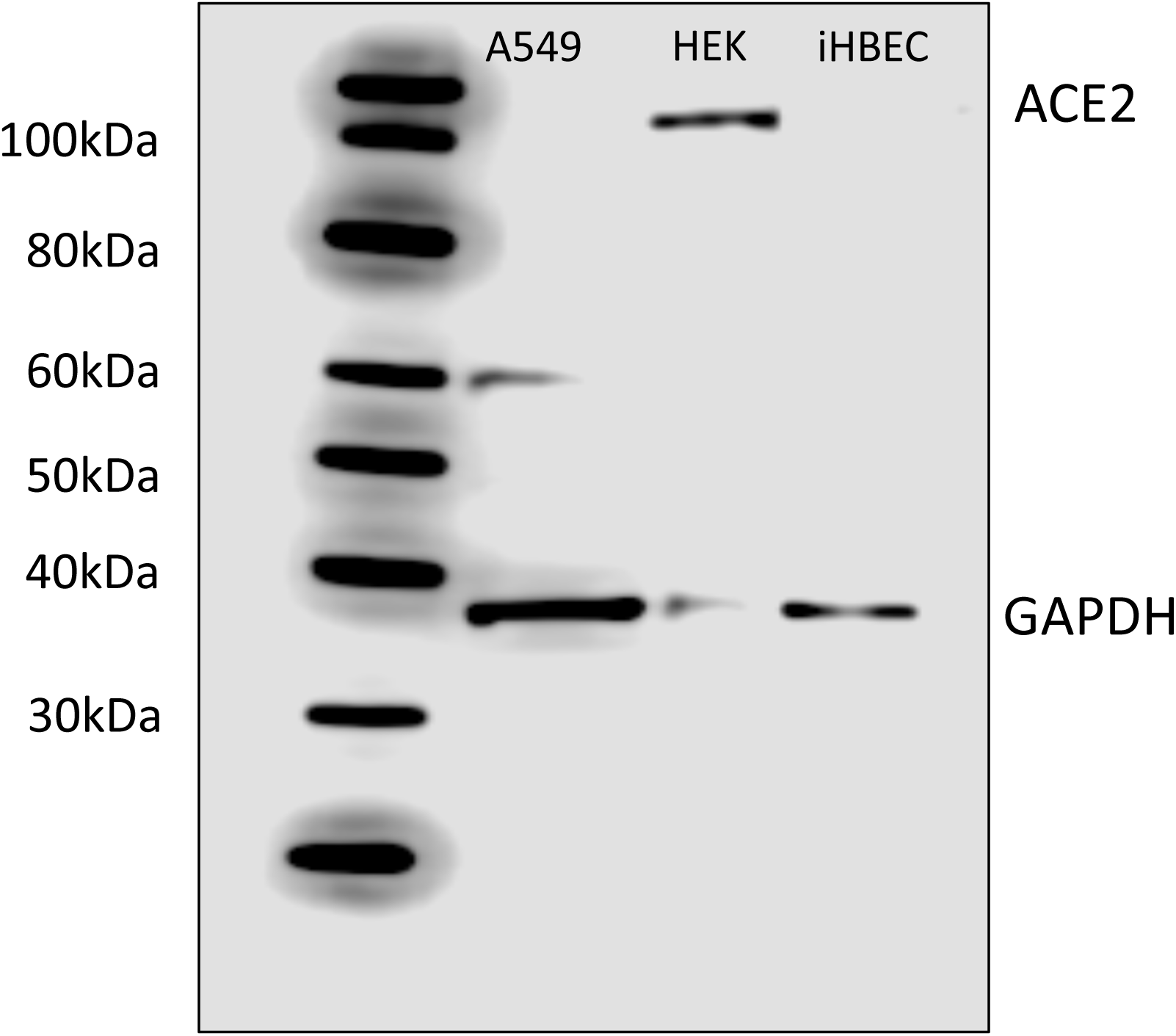
Independent confirmation of immunoblot analysis for ACE2. Lane 1 = A549 cell line. Lane 2 - HEK293 cells. Lane 3 - immortalized human bronchial epithelial cells. ACE2 has a predicted molecular weight of 110kDa with GAPDH as a loading control. Anti-human ACE2 antibody is distinct from immunoblot in Figure 4.

**Supplement Figure 3:**
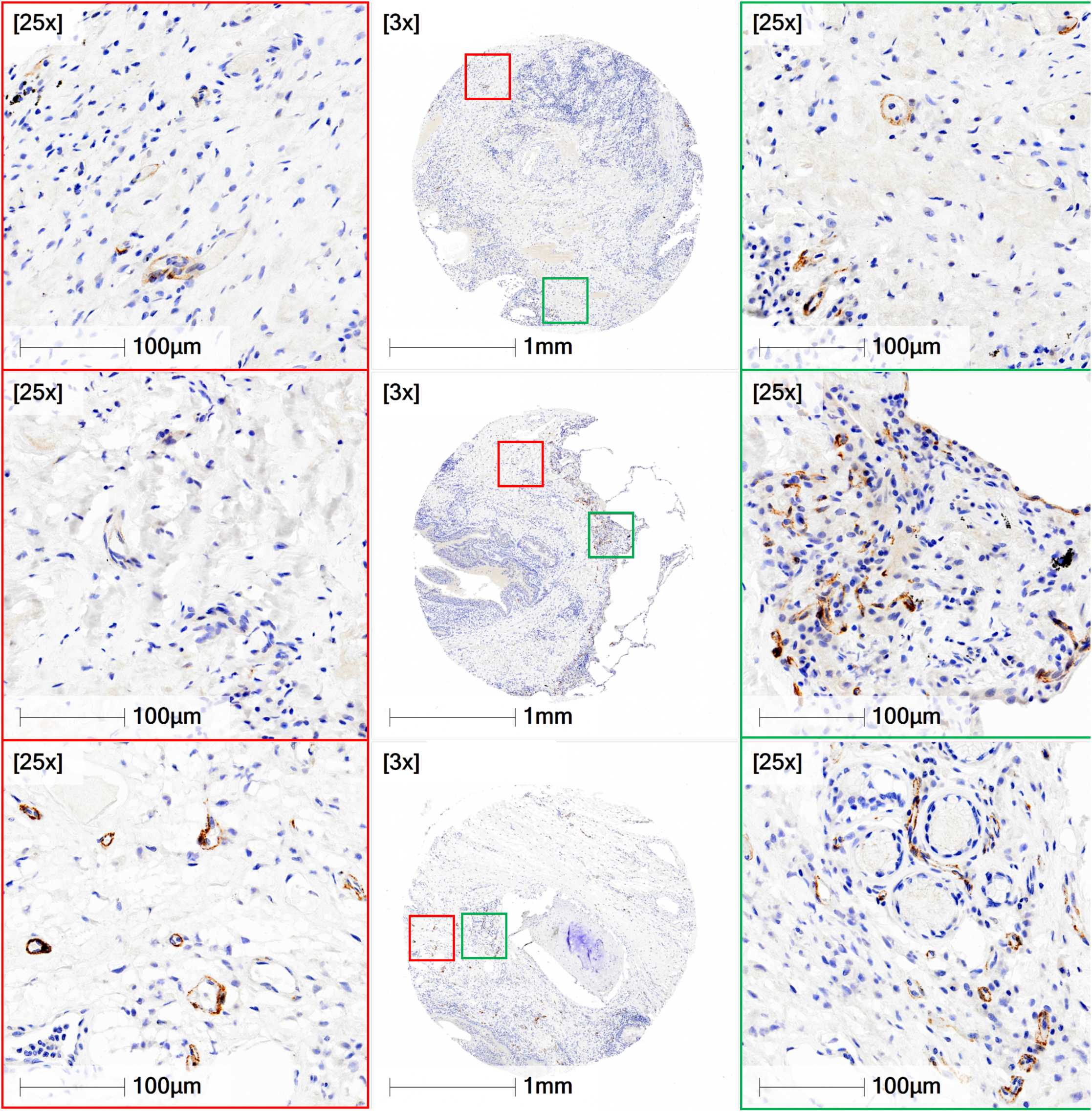
Immunohistochemical localization of ACE2 in microvasculature of human lung tissue. Representative examples (n=3 donors) of positive ACE2 protein staining (rust/brown) in human lung tissue in regions distinct from those fields of view containing conducting airways. Images taken from identical slide use for Figure 5 (same staining run and conditions for image acquisition). Red and green boxes are 60X zoom of 3X magnification of entire tissue core sample.

**Supplement Figure 4:**
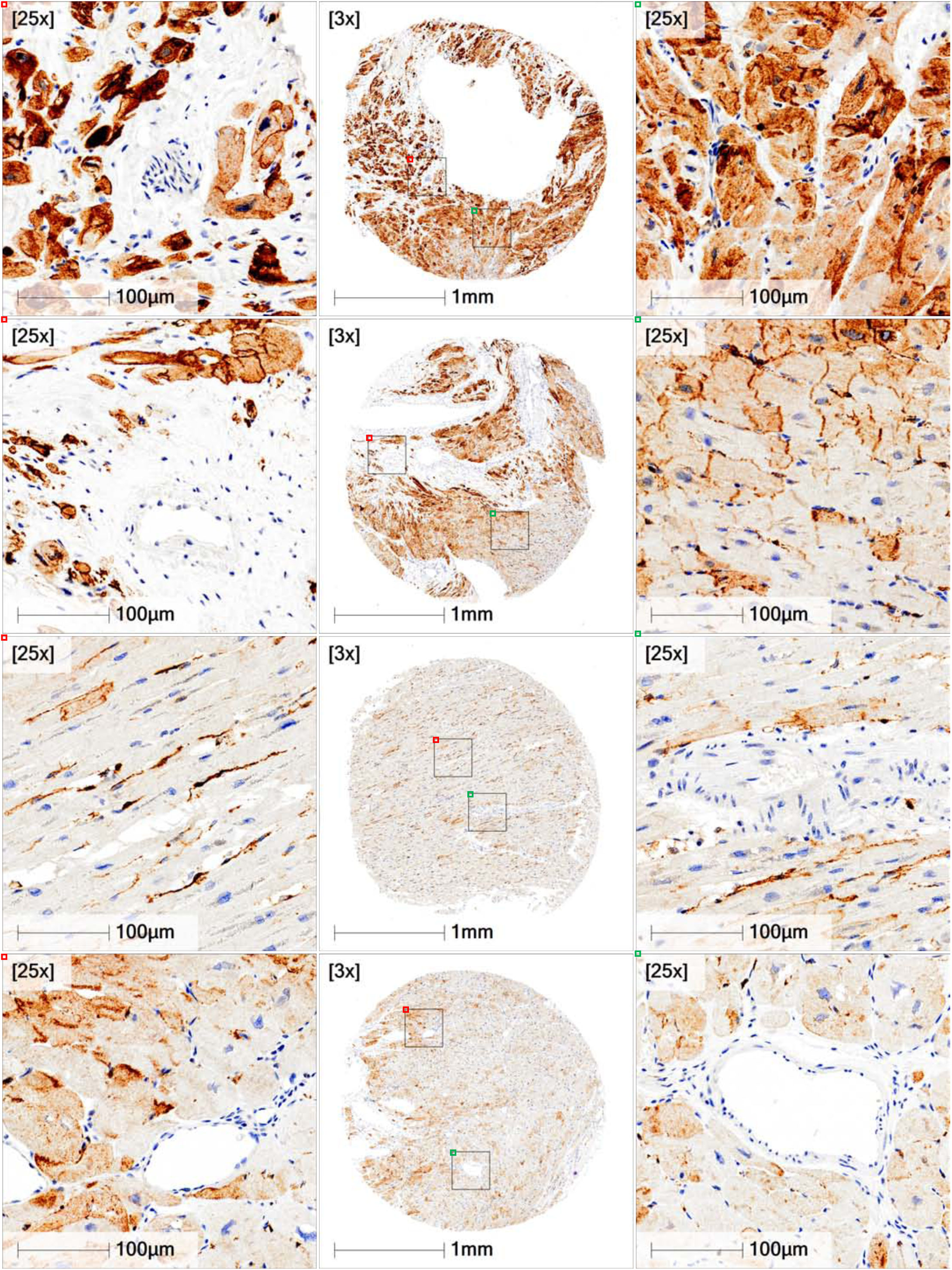
Immunohistochemical localization of ACE2 in human heart tissue. Representative examples (n=4 donors) of positive ACE2 protein staining (rust/brown) in human heart tissue. Staining protocol identical to Figure 5 and Supplementary Figure 3. Heart tissues stained on same staining run on Leica Bond Rx autostainer as for lung tissue in Figure 5. Red and green boxes are 60X zoom of 3X magnification of entire tissue core sample.

